# Sensitivity to Monoclonal Antibody 447-52D and an Open Env Trimer Conformation Correlate Poorly with Inhibition of HIV-1 Infectivity by SERINC5

**DOI:** 10.1101/2020.04.10.036483

**Authors:** Aaron O. Angerstein, Charlotte A. Stoneham, Peter W. Ramirez, John C. Guatelli, Thomas Vollbrecht

## Abstract

The host protein SERINC5 inhibits the infectivity of HIV-1 virions in an Env-dependent manner and is counteracted by Nef. The conformation of the Env trimer reportedly correlates with sensitivity to SERINC5. Here, we tested the hypothesis that the “open” conformation of the Env trimer revealed by sensitivity to the V3-loop specific antibody 447-52D directly correlates with sensitivity to SERINC5. Of five Envs tested, SF162 was the most sensitive to neutralization by 447-52D, but it was not the most sensitive to SERINC5; instead the Env of LAI was substantially more sensitive to SERINC5 than all the other Envs. Mutational opening of the trimer by substitution of two tyrosines that mediate interaction between the V2 and V3 loops sensitized the Envs of JRFL and LAI to 447-52D as previously reported, but only BaL was sensitized to SERINC5. These data suggest that trimer “openness” is not sufficient for sensitivity to SERINC5.

## Introduction

The HIV-1 envelope glycoprotein (Env) mediates viral infectivity. Proteolytic cleavage of the gp160 precursor protein leads to the formation of Env spikes comprising trimeric heterodimers of gp120 and gp41. The Env spike protrudes through viral and cellular membranes and is responsible for the binding to and the fusion of infectious virus particles (virions) with target cells (Berger et al., 1999).

The structure of the Env trimer transitions between at least three conformations – closed, partially open, or open – although one conformation tends to predominate in a given Env (Cai et al., 2017; Lu et al., 2019; Ma et al., 2018; Montefiori et al., 2018). Envs whose trimer conformation is open are usually found in laboratory-adapted strains that are easily neutralized by many antibodies. On the other hand, primary Envs derived directly from HIV-infected patients typically have a relatively closed trimer conformation, and the epitopes recognized by some neutralizing antibodies are not accessible (Brandenberg et al., 2015; Cai et al., 2017). For example, the antibody 447-52D, which recognizes an epitope at the apex of the V3 loop (Stanfield et al., 2004), neutralizes the open trimers of so-called tier 1 Envs, whereas, the more closed trimers of tier 2 and 3 Envs, which include the majority of Envs from primary isolates, are resistant to the antibody 447-52D (Seaman et al., 2010).

The HIV-1 accessory protein Nef is a small peripheral membrane protein that plays several roles in the viral pathogenesis, one of which is the enhancement of infectivity (Chowers et al., 1994; Deacon et al., 1995; Kestler et al., 1991). Nef downregulates cell surface host proteins such as CD4 and the major histocompatibility complex class-I (MHC-I), but the enhancement of infectivity is due, at least in some cell lines and in primary CD4-positive T lymphocytes, to the removal of SERINC5 from the plasma membrane (Rosa et al., 2015; Usami et al., 2015). In the absence of Nef, SERINC5 incorporates into virions and inhibits the infectivity of cell-free virus.

The magnitude of the *nef*-phenotype varies in an *env*-dependent manner; this was ultimately explained by the observation that Env proteins are differentially sensitive to SERINC5. First, the Env of JRFL was noted to render pseudovirions insensitive to Nef (Lai et al., 2011). The difference between the Env of JRFL, which does not support a Nef-effect, and the Env of SF162, which does, was then mapped to the variable regions of Env, specifically the so-called B-C loop in V2 (Usami and Göttlinger, 2013). Moreover, whether or not these Env proteins supported a Nef-effect correlated with sensitivity to neutralization by the antibody 447-52D. These data led to the conclusion that the enhancement of infectivity by Nef depended on an open quaternary conformation of the Env trimer (Usami and Göttlinger, 2013). The last piece of this puzzle fell into place with the discovery of SERINC5 as an inhibitor of infectivity counteracted by Nef. The Env of SF162 was sensitive to SERINC5, whereas the Env of JRFL was not, and this difference mapped to the same variable region of Env that determined Nef-responsiveness and had been shown to affect sensitivity to the antibody 447-52D (Usami et al., 2015).

These findings suggested that the majority of Envs in primary HIV-1 isolates or clones, whose trimers are more closed, might be resistant to SERINC5; this possibility was supported experimentally (Beitari et al., 2017). On the other hand, CD4, if unopposed by Nef, seemed able to sensitize Envs that would otherwise appear resistant to SERINC5 (Zhang et al., 2019).

We aimed to test the hypothesis further that Env trimer conformational openness is a determinant of sensitivity to SERINC5. To do this, we evaluated five Env proteins (SF162, JRFL, LAI, AC10, and 1012)-two of which we expected to be relatively sensitive to SERINC5 (LAI and SF162) and three of which we expected to be resistant (JRFL, AC10, and 1012) - for sensitivity to the antibody 447-52D and to the expression of SERINC5, each in a dose-response format. We also used specific mutations in the V2 loop of Env to open the trimer, both in the context of Env from JRFL (relatively resistant to SERINC5) and from BaL (which we found to be relatively sensitive to SERINC5). Our data suggest that trimer openness is likely necessary for, but not the only determinant of, SERINC5-sensitivity.

## Results

### Selection of Env proteins for comparison

HIV-1 Envs can be classified based on their neutralization “tier”: tier 1A Envs are highly sensitive to neutralization whereas tiers 2 and 3 are relatively resistant to neutralization. A relationship between neutralization-sensitivity and Env-trimer-conformation is apparent: open in the case of tier 1 Env trimers versus closed in case of tier 3 Envs (reviewed in Montefiori et al., 2018). Here, we aimed to test and extend the hypothesis that open trimers identified by sensitivity to the antibody 447-52D were relatively sensitive to SERINC5, whereas the more closed trimers identified by resistance to 447-52D were relatively resistant to SERINC5. We chose the following Envs for initial comparison: SF162, JRFL, LAI, AC10, and 1012. The Envs SF162, AC10 and 1012 have been previously characterized for their sensitivity to 447-52D (Seaman et al., 2010): SF162 is highly sensitive and classified as tier 1A, whereas 1012 and AC10 are resistant and classified as tier 1B and tier 2, respectively (Seaman et al., 2010). The Env of JRFL was characterized previously as resistant to 447-52D (Guzzo et al., 2018; Usami and Göttlinger, 2013), whereas data indicating the sensitivity of the Env of LAI are lacking. Regarding sensitivity to SERINC5, the Env of SF162 has previously been characterized as sensitive, whereas that of JRFL has been characterized as resistant (Usami 2013). We expected the Env of LAI to be sensitive to SERINC5, since it is closely related to the Envs of NL4-3 and HXB2, with which the influence of Nef on infectivity was discovered (Chowers et al., 1994; Miller et al., 1994). The sensitivity of the Envs AC10 and 1012 to SERINC5 were not previously known.

### Infectivity of pseudovirions containing the various Envs and their sensitivities to 447-52D and SERINC5

We generated HIV-1 pseudoviruses using HEK 293T cells that were co-transfected with three plasmids: an NL4-3-based ΔEnv-ΔNef proviral plasmid, an HA-tagged SERINC5 expression plasmid, and a plasmid expressing the indicated Env. The pseudoviruses were collected after 72 hours after transfection, partially purified, and used to infect TZM-bl reporter cells, in which luciferase activity is expressed under the control of the HIV-1 LTR and activated by the expression of Tat. Luciferase activity (relative light units, RLUs) was measured in the lysates of the TZM-bl cells 48 hours after infection. The infectivity of the various preparations of pseudovirions (Fig. 1A) was variable between quadruplicate, independent experiments (Figure 1A). Consequently, in the following inhibition experiments, the data are shown for each preparation relative to its infectivity in the absence of the respective inhibitor (antibody 447-52D or SERINC5).

**Figure 1.**
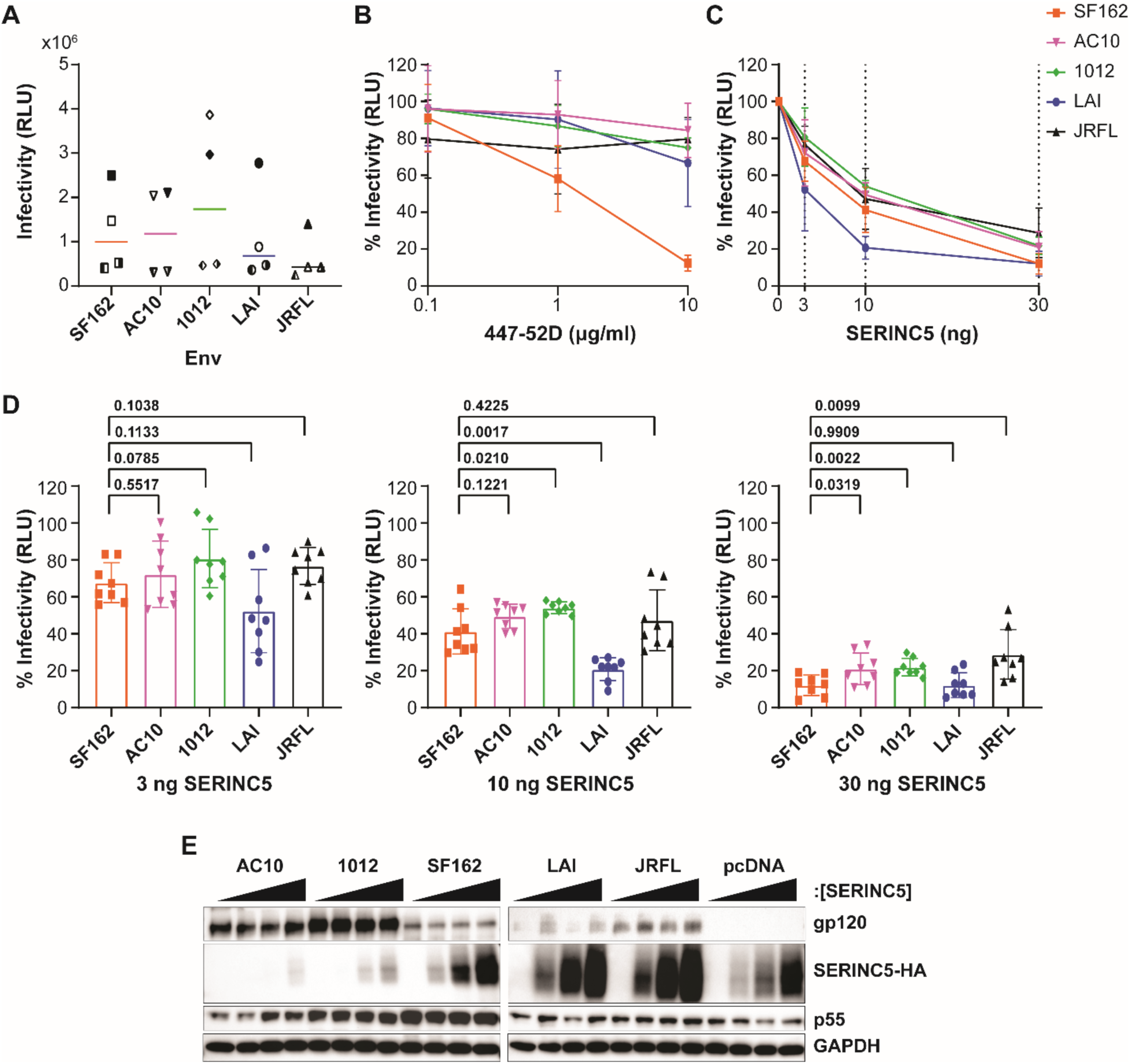
Relative sensitivity of different Envs to 447-52D antibody and SERINC5. A) Pseudovirions comprising DHIV-ΔEnv-ΔNef and Env SF162, AC10, 1012, LAI or JRFL were produced in HEK 293T cells and infectivity measured by in HeLa TZM-bl luminescent reporter cells, as described in the Methods section. The relative light unit (RLU) values for four different preparations of pseudovirion stocks are shown for each Env; each preparation was measured in triplicate, and the mean of the four independent experiments is shown for each Env. Experiment 1 – filled symbol; 2 – the left half of symbol filled; 3 – the right half of symbol filled; 4 – symbol blank. B) Neutralization of infectivity by the antibody 447-52D. The relative infectivity of pseudovirions is shown on the y-axis after normalization to no antibody added control of the indicated Envs (0 μg/ml 447-52D = 100%), at increasing antibody concentrations from 0.10 μg/ml to 10 μg/ml. The data shown represent two independent experiments; with each neutralization assay performed in duplicate. C) SERINC5-mediated inhibition of infectivity. The relative infectivity of pseudovirions comprising indicated Envs is shown, following normalization to the no-SERINC5 plasmid control for each of the indicated Envs (0 ng SERINC5 = 100%). The dashed lines indicate the amount of SERINC5 plasmid used (0, 3, 10, or 30 ng). The data represents four independent experiments (pseudovirion preparations); each data point represents the mean of eight independent values for each Env (the infectivity of each pseudovirion preparation measured at 1:3 and 1:9 dilutions, each in triplicate). D) Relative infectivity of pseudovirions at increasing concentrations of SERINC5. Independent infectivity data values are compared for each concentration of SERINC5 for experiments presented in panel C. Pairwise comparisons were made between each Env to SF162 using Welch’s t-test (p values are shown). E) Expression of Env and SERINC5 in the HEK 293T virus producer cells. The HEK 293T producer cells were lysed and proteins separated by SDS-PAGE followed by immunoblotting for the indicated proteins. Each Env was grouped by increasing SERINC5 protein concentration: 0, 3, 10, then 30 ng from left to right. The different Envs are indicated; pcDNA is a no-Env control.

The sensitivity of each Env to the monoclonal antibody 447-52D was tested in duplicate in two independent neutralization assays (Figure 1B). SF162 was sensitive to the 447-52D, whereas JRFL, AC10, and 1012 were resistant, in agreement with previous data. Unexpectedly, LAI was relatively resistant to the antibody 447-52D and similar to AC10, 1012, and JRFL. Sequencing of the LAI expression-plasmid confirmed that the linear epitope at the apex of the V3 loop recognized by 447-52D was present (data not shown), leaving the basis for its resistant phenotype unexplained.

Next, we assessed the sensitivity of each Env to increasing concentrations of SERINC5 using four independent preparations of pseudovirions (Figure 1, C and D). Figure 1C indicates that each Env was sensitive to the expression of SERINC5 in a dose-dependent manner. Notably, the Env of LAI appeared substantially more sensitive to SERINC5 than the other Envs, which were similar to each other. To more carefully compare the Envs with statistical analyses, we represented the data of Figure 1C as bar graphs showing the infectivity relative to no expressed SERINC5 for the 3, 10, and 30 ng amounts of transfected SERINC5 plasmid (Figure 1D); statistical significance was assessed for the differences between each Env compared to SF162, since SF162 was the only Env that was markedly sensitive to 447-52D. Differences between the Envs were of very small effect-size; nonetheless, when 10 ng of SERINC5-plasmid was used, 1012 appeared less sensitive to SERINC5 than SF162, whereas LAI appeared more sensitive; when 30 ng of SERINC5-plasmid was used, 1012, AC10, and JRFL all appeared less sensitive to SERINC5 than SF162.

We assessed protein expression in the transfected HEK 293T cells that produced the pseudovirions for one of the replicate experiments whose data are included above (Figure 1E). The western blots confirmed similar expression of p55 across the SERINC5-plasmid dose ranges for each Env. They confirmed the expression of Env proteins, although the strength of the signals varied, potentially due to actual differences in Env expression or differential efficiency of detection by the polyclonal antibody to gp120. The expression of SERINC5 increased as expected, with an increasing amount of transfected plasmid. However, in this and other replicate experiments (data not shown), the expression of SERINC5 seemed affected by Env: relative to no-Env (“pcDNA”), the expression of SERINC5 seemed reduced when AC10 and 1012 were expressed, whereas it seemed increased when LAI and JRFL were expressed.

### Mutational opening of the Env trimer using mutations in the V2 loop

As an alternative to the approach of comparing different Envs, we asked whether opening the trimer mutationally could sensitize Env to SERINC5. We took advantage of previous data indicating that substitution of the tyrosines 173 and 177 in V2 with phenylalanine increases sensitivity to various agents consistent with trimer opening including the 447-52D antibody (Guzzo et al., 2018). The substitution of these tyrosines was proposed to disrupt the interaction of V2 with V3, thereby opening the trimer. Here, we studied these substitutions in the context of two Envs: JRFL and BaL (Guzzo et al., 2018). As noted above, JRFL - a tier 2 Env - is reportedly resistant to 447-52D and relatively resistant to SERINC5, whereas BaL - a tier 1B Env - is sensitive to 447-52D (Guzzo et al., 2018; Seaman et al., 2010). The sensitivity of BaL to SERINC5 was not previously known.

As above, replicate preparations of pseudovirions were variable in infectivity, but on average, the phenylalanine substitution mutants (“FF”) reduced the infectivity of both JRFL and BaL Envs (Figure 2A). As expected, based on the published data (Guzzo et al., 2018), BaL was more sensitive to 447-52D than JRFL, and the FF substitution sensitized both Envs to the antibody (Fig. 2B). The BaL Env was more sensitive to SERINC5 compared to JRFL (Figure 2C), consistent with the hypothesis that lower-tier Envs with more open trimers are more sensitive to SERINC5. The FF substitution modestly sensitized BaL to SERINC5, but it did not detectably sensitize JRFL to SERINC5 (Figure 2C). As above, we re-graphed the data for each amount of SERINC5 plasmid and analyzed the comparisons between the wild type Envs and their phenylalanine substitution counterparts (Figure 2D), this analysis indicated that low p values were present only at the 30 ng amount of SERINC5-plasmid. Under this condition, the sensitivity of BaL to SERINC5 was increased by the phenylalanine substitutions, but the sensitivity of JRFL to SERINC5 was not. A western blot of the pseudovirion-producer cells from one of the experiments used to generate the data of Figure 2C and D is shown in Figure 2E; this blot confirmed the appropriate expression of p55 and Env and dose-dependent expression of SERINC5.

**Figure 2.**
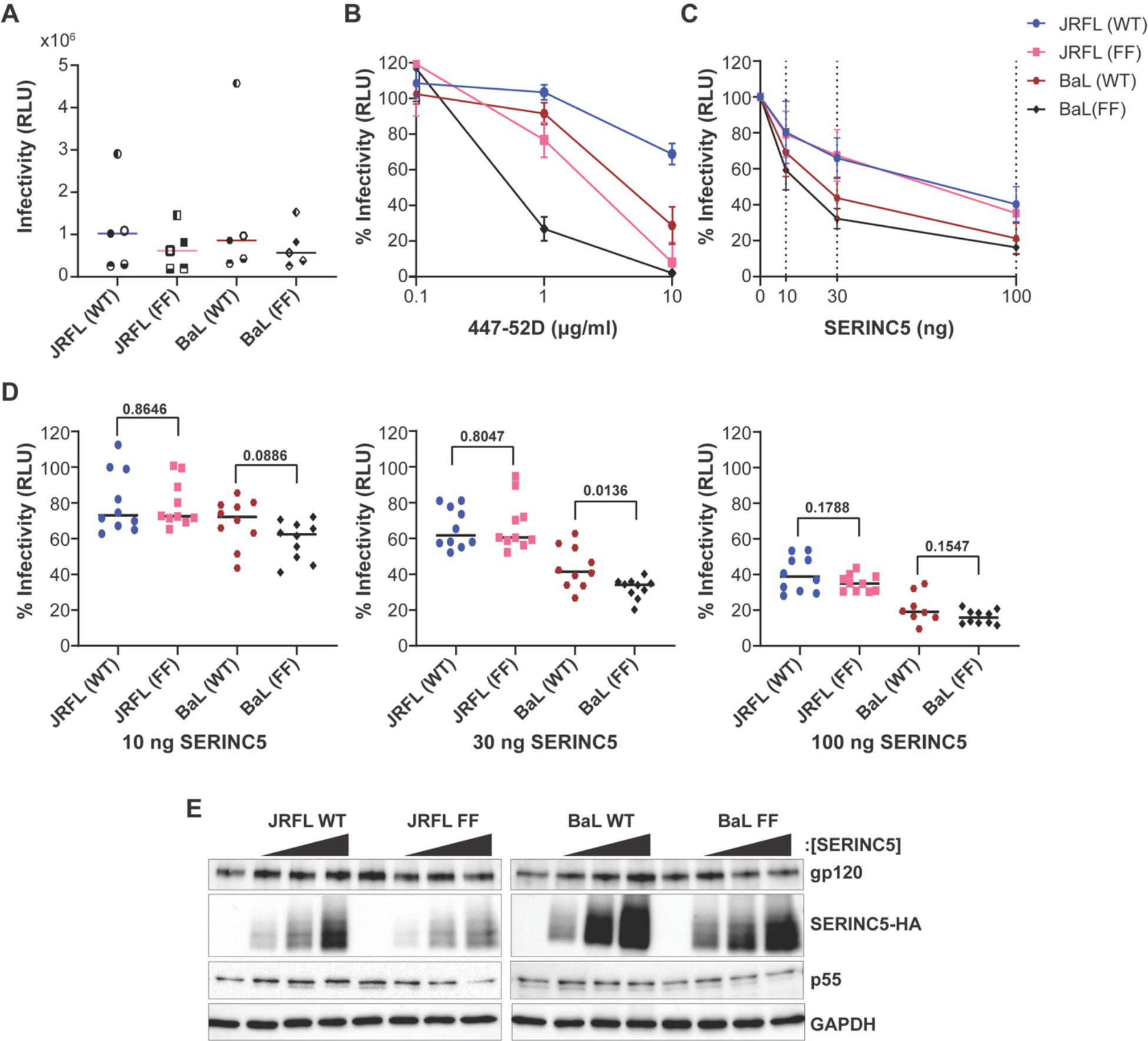
Effect of V2 tyrosine substitutions on the sensitivity 447-52D antibody and to SERINC5. A) The infectivity of pseudovirions comprising WT or mutant “FF” (YY173, 177 FF) Envs JRFL or BaL is shown in Figure 1A. The means from five independent experiments are shown for each Env; and each assay was run in triplicate. Experiment 1 – filled symbol; 2 – symbol blank; 3 – the top half of symbol filled; 4 – the bottom half of symbol filled; 5 – left half of the symbol filled. B) Neutralization of infectivity by the antibody 447-52D. Data are presented as in Figure 1B. The data is shown for two independent experiments; each neutralization assay was run in duplicate. C) Inhibition of infectivity by expression of SERINC5. The relative infectivity of pseudovirions was quantified each of the indicated Envs and compared to no added SERINC5 (0 ng SERINC5 = 100%). The dashed lines indicate the amount of SERINC5 added (0, 3, 10, and 30 ng). Data are shown from five independent experiments; each data point represents 10 independent values for each Env (the infectivity of each pseudovirion preparation measured at 1:3 and 1:9 dilutions, each in triplicate). D) Relative infectivity of pseudovirions comprising WT and mutated Envs at increasing concentrations of SERINC5. Pairwise comparisons were made between each Env and its corresponding tyrosine-to-phenylalanine mutant. P-values were determined by Welch’s t-test. E) Expression of the key proteins was measured in the pseudovirion-producer cells. Molecular masses in kDa are indicated on the left; Proteins are indicated on the right. Each Env was grouped by increasing SERINC5 protein concentration: 0, 3, 10, then 30 ng from left to right.

## Discussion

We sought to test and extend the hypothesis that the “openness” of the Env trimer is the key determinant of sensitivity to the host protein SERINC5. We used two approaches: comparison of different Envs of different tiers, and mutational opening of the trimer to allow comparison with the same Env. In both cases, we used sensitivity to the V3 loop antibody 447-52D as a measure of trimer openness. The comparison of different Envs was fraught with an unexpected paradox: the Env of LAI was relatively resistant to the monoclonal antibody 447-52D, despite the fact that it was more sensitive to SERINC5 than any of the other Envs. Moreover, while the group of Envs resistant to 447-52D (JRFL, AC10, and 1012) were clearly more resistant to SERINC5 than LAI, they were only slightly more resistant to SERINC5 compared to SF162, which was highly sensitive to 447-52D. Indeed, a remarkable conclusion of our study is that, in the absence of Nef and under the conditions of SERINC5 expression herein, all the Envs tested were sensitive to SERINC5. Similarly, the mutational opening of the trimer did not support that this quality of Env is the sole determinant of sensitivity to SERINC5: disruption of V2-V3 interaction by substitution of tyrosines in V2 sensitized the Envs of BaL and JRFL to 447-52D as previously reported, but sensitization to SERINC5 was minimal (BaL) to none (JRFL). Overall, these data suggest that while trimer openness might correlate with SERINC5-sensitivity at some level, it is not the sole or simple determinant.

Our study has several limitations and caveats. The various Envs tested here likely have many differences in addition to trimer openness that might affect sensitivity to SERINC5. Moreover, sensitivity to the antibody 447-52D might not be a perfect surrogate for trimer openness. The latter is potentially exemplified by LAI, which we expected to behave as a tier 1 Env - sensitive to 447-52D; it was instead relatively resistant, and this resistance was not explained by mutation of the epitope itself (data not shown). Lastly, we expressed SERINC5 by transient transfection. This allowed for the rigor of the dose-response experimental design, but despite our use of low amounts of plasmid, SERINC5 is potentially over-expressed here relative to cells that naturally host HIV-infection.

Our study also has an additional anomalous observation: the expression of SERINC5 seemed to be affected differently by the various Env constructs. Although this observation must be considered at best highly preliminary, it is intriguing insofar as an effect of Env on SERINC5 expression could be a mechanism in addition to trimer openness by which the apparent sensitivity to Env could vary.

Insofar as trimer openness is a determinant of SERINC5-sensitivity, we note that the tyrosine substitutions utilized here to open the trimer (Guzzo et al., 2018) are within the V2 region previously identified as a determinant of the differential sensitivity of JRFL and SF162 Envs to SERINC5 (Usami et al., 2015). While we did not confirm the results of domain swaps between JRFL and SF162, the tyrosine substitutions in V2, as shown here, did not sensitize the JRFL Env to SERINC5, despite that it sensitized this Env to 447-52D as reported.

In summary, our data suggest that trimer openness as assessed by sensitivity to the antibody 447-52D is not the sole determinant of the sensitivity of Env proteins to inhibition by SERINC5. The Env of LAI appears especially sensitive to SERINC5 for unknown reasons. Moreover, although our study is a small sample, both tier 1 and 2 Envs appear to be sensitive to SERINC5 under the conditions tested.

## Materials and Methods

### Molecular Clones

Three molecular clones expressing Env were obtained through the NIH AIDS Reagents Program, Division of AIDS, NIAID, NIH: HIV-1 SF162 gp160 Expression Vector (pCAGGS SF162) from Drs. L. Stamatatos and C. Cheng-Mayer (Cheng-Mayer et al., 1997; Stamatatos et al., 2000, 1998), HIV-1 1012 Env Expression Vector (p1012.TC21.3257) from Drs. Beatrice H. Hahn, Brandon F. Keele, and George M. Shaw (Keele et al., 2008), and HIV-1 AC10.29 Env Expression Vector (AC10.0, clone 29 (SVPB13)) from Dr. David Montefiori and Dr. Feng Gao (Li et al., 2005). pLET-LAI, pLET JRFL and DHIV plasmids were a kind gift from Vicente Planelles (Bonczkowski et al., 2014; Bosque and Planelles, 2009; Challita-Eid et al., 1998). The pCAGGS-JRFL and pcDNA(3)-BaL.01 were a kind gift from Paulo Lusso (Guzzo et al., 2018). All clones were verified by sequencing, and the plasmid DNA was purified using the Wizard Plus SV Minipreps DNA Purification System (Promega) or QIAGEN Plasmid Midi kits.

### Cell Culture

HEK 293T cells and TZM-bl cells were cultured in Dulbecco’s Modified Eagle’s Medium (DMEM - Gibco) supplemented with 10% fetal bovine serum (FBS) and 100 U/ml penicillin-streptomycin (Gibco); cells were incubated at 37ºC in 5% CO_2_. The following reagent was obtained through the NIH AIDS Reagent Program, Division of AIDS, NIAID, NIH: TZM-bl cells (Cat# 8129) from Dr. John C. Kappes, and Dr. Xiaoyun Wu (Derdeyn et al., 2000; Platt et al., 2009, 1998; Takeuchi et al., 2008; Wei et al., 2002).

### Transfection

HEK 293T cells were seeded in 12-well plates at a density of 1.5 × 10^5^ cells in one ml of complete media per well. The cells were transfected using Lipofectamine 3000 (Thermo Fisher Scientific), according to the manufacturer’s instructions; 625 ng of DHIV-∆Env-ΔNef, and 375 ng of the desired Env expression plasmid were mixed with 0, 3, 10, 30, or 100 ng pBJ5-SERINC5-HA or empty pBJ5 plasmid and Opti-MEM media (Gibco). The transfection mix was then added to Lipofectamine 3000 diluted in Opti-MEM and incubated at room temperature for 15 minutes. 100 μl of the transfection mix was added per well. The plates were then incubated for 72 hours at 37°C.

### Infectivity Assay

Supernatant was collected from each well and centrifuged for five minutes at 300 × g to remove cells and debris. Virions were harvested by centrifugation of culture supernatants through a 20% sucrose cushion at 23,500 × g for one hour at 4°C. The pelleted pseudovirions were resuspended in 600 μl of complete media and dilutions (1:3 and 1:9) were made for each pseudovirion-preparation. The diluted virions were mixed with 2 × 10^4^ TZM-bl cells in 100 μl DMEM per well in 96-well black-wall plates (Corning), in triplicate. The cells were incubated for 48 hours at 37°C. The supernatant was then removed, and cells lysed in 70 μl of 0.5% Triton-X, before addition of 70 μl of Britelite Plus Luciferase reagent (Perkin Elmer). Luminescence was quantified using a Perkin Elmer luminometer. Data are presented as relative light units (RLUs) following subtraction of background controls.

### Neutralization Assay

We followed the protocol for neutralizing antibody assays provided by the Montefiori Laboratory at Duke University (Montefiori, 2009). In each well of a 96-well black-wall plate, 45 μl pseudovirions were combined with 5 μl of 447-52D antibody dilutions, in duplicate. The plate was then incubated for 90 minutes at 37°C before addition of 1 × 10^4^ TZM-bl cells in 100 μl of complete DMEM per well. The plates were incubated for 48 hours at 37°C and CO_2_ before quantification of cellular luciferase activity, as described above. The following reagent was obtained through the NIH AIDS Reagent Program, Division of AIDS, NIAID, NIH: Anti-HIV-1 gp120 Monoclonal (447-52D) from Dr. Susan Zolla-Pazner (Gorny et al., 1992).

### Western Blots

The HEK 293T virus producer cells were briefly washed with PBS and resuspended using Acutase (Corning) cell suspension solution. The cells were pelleted by centrifugation at 300 × g for 5 minutes, resuspended in ice-cold lysis buffer (50 mM NaCl, 1% Triton-X-100, 50 mM Tris, pH 8.0) supplemented with cOmplete^TM^ protease inhibitor cocktail (Roche) and agitated for 30 minutes at 4°C. The nuclei were pelleted by centrifugation at 13,200 × g for 20 minutes at 4°C. The protein concentration of the supernatant was determined by Quick Start ^TM^ Bradford assay (Bio-Rad) following standard protocol, and equal protein concentrations diluted in 2x Laemmli sample buffer containing 100 mM tris (2-carboxyethyl) phosphine hydrochlorine (TCEP-HCl) reducing agent. Protein samples and a molecular weight size marker (Pageruler Plus, ThermoFisher Scientific) were separated on 10% SDS-PAGE gels and transferred to PVDF membrane. The membranes were blocked in 5% skim milk diluted in Phosphate-Buffered Saline with 0.2% Tween-20 (PBS-T) for one hour at RT, and incubated with the following primary antibodies (diluted in 1% skim milk-PBS-T) overnight at 4°C; Envelope was detected with sheep α-gp120 (NIH #288, 1:5000), SERINC5-iHA was detected with mouse α-HA (clone 16B12, Biolegend, diluted 1:5000), Gag was detected using mouse α-p24 (Millipore, diluted 1:5000), and GAPDH was detected with mouse α-GAPDH (GeneTex, diluted 1:5000). The membranes were washed extensively with PBS-T before incubated with goat anti-mouse-HRP (Bio-Rad Laboratories) or rabbit anti-sheep-HRP (Bio-Rad Laboratories) secondary antibodies (diluted 1:3000). The blots were washed with PBS-T and incubated with Clarity™ Western ECL Blotting reagent (Bio-Rad Laboratories). Chemiluminescence was detected using the ChemiDoc™ Imaging System and analyzed using Image Lab software v5.1 (Bio-Rad Laboratories).

The following reagent was obtained through the NIH AIDS Reagent Program, Division of AIDS, NIAID, NIH: Antiserum to HIV-1 gp120 from Dr. Michael Phelan (Hatch et al., 1992; Page et al., 1992).

### Statistical Analysis

Normalization of infectivity assay data to the no-SERINC5-plasmid or no-antibody controls and the generation of graphs and Welch’s t-test were done using GraphPad Prism 8. Figures were produced using Adobe Creative Cloud v2020 Photoshop and Illustrator software.

## Acknowledgments

We thank Dr. Heinrich Göttlinger for the pBJ5-SERINC5-iHA expression plasmid; Dr. Vicente Planelles for the pNL43-derived ΔEnv-ΔNef (DHIV-ΔNef), pLET-LAI and pLET-JRFL plasmids; and Dr. Paolo Lusso for the expression plasmids for JRFL and BaL and their isogenic tyrosine to phenylalanine V2 mutants. This work was supported by NIH NIAID R01AI129706 to JG.

